# *In vitro* evaluation of *Escherichia coli* and *Staphylococcus aureus* translocation in 3D printed material

**DOI:** 10.64898/2026.02.02.703277

**Authors:** Ashma Sharma, Joshua Prince, A-Andrew D Jones

## Abstract

Vascular graft infection is a rare but life threating condition, primarily occurring after 30 days post-surgery. Meta-analysis has shown that antimicrobial coatings on graft materials do not prevent these infections. Moreover, infection still occurs even though studies have also shown that there is no bacterial proliferation on or bacterial penetration of common vascular graft material. The time frame of infection, meta-analysis, and *in situ* studies suggest that bacteria present at the suture site are introduced into the surrounding tissue or that systemically circulating bacteria may be surviving, proliferating, diffusing slowly, and evading host immune defense in synthetic vascular grafts. *De novo* vascular graft materials, such as tissue-engineered vascular graft material and decellularized vasculature may provide an *in situ* platform for studying survival, proliferation, and diffusion in tissue and tissue-like materials. In this study, we use confocal microscopy to image penetration depth of bacteria over time as a proxy for diffusion of *Staphylococcus aureus* and *Escherichia coli* into alginate, GelMA, and decellularized porcine vascular tissue. We quantified viable bacteria breakthrough as a function of biomaterial type. We found penetration depth over time was similar in all three biomaterials, however *E. coli* broke through much less from tissue than from engineered materials, while *S. aureus* had higher breakthrough in the GelMa but otherwise equal rates. These results point to the possibility of interstitial growth control relative to surface coatings as a future target for engineering infection resistance in engineered vascular grafts.

**Graphical Abstract:** 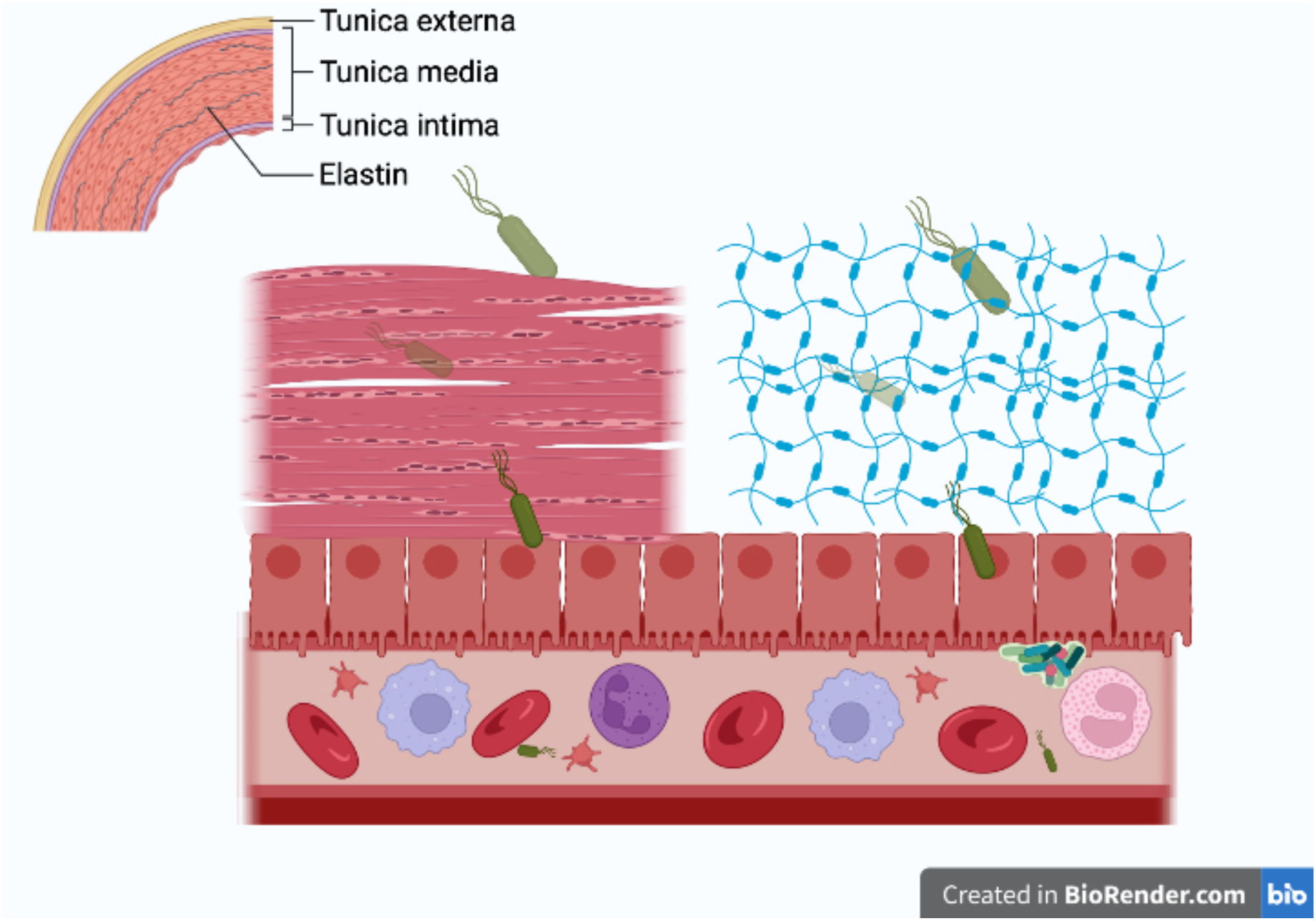

## Introduction

Vascular grafting has been used in coronary artery bypass surgeries and to treat peripheral artery disease trauma^1-6^. These surgeries are common features of care for the 607 million globally suffering from cardiovascular disease; the 237 million globally suffering from peripheral artery disease and trauma, as well as others suffering from rare, but often deadly, congenital heart defects and Kawasaki Disease^1^. Autologous vascular grafts, like the saphenous vein for coronary artery bypass surgery, provide the necessary vessels for most patients^1,7-9^.

Engineered vascular grafts are viewed as less invasive than autologous vascular grafts with higher long-term survival rates than non-surgical interventions like stents ^7-9^. Engineered vascular grafts are used for patients at high risk for complications in successive surgeries, such as harvesting vessels, for small vessels (<6 mm ID), and for others that lack healthy vessels to harvest^10,11^. Despite wide availability and biocompatibility, engineered vessels maintain patency at lower rates, occlude more rapidly, and exhibit higher rates of infection than autologous vascular grafts. The burden of these failures falls particularly on women, who receive more engineered vascular grafts to treat peripheral artery disease than men^12^. While there has been progress in engineered vascular grafts, including small diameter grafts, there is little data on the rate and type of infections of engineered vascular grafts^13-17^.

Vascular graft infections occur more often in later stages (> 30 days) after surgery^18^. Overall, vascular graft infection is rare, occurring for example in 1-3% of surgeries in Switzerland in 2013 ^18^. However, mortality rates from infected vascular grafts can range from nearly 20% over one year to 40% over four years with a financial cost of ∼$80,000 per patient in 2013 ^18^. This high mortality rate is generally assumed to be due to the breakdown of engineered vascular grafts and is one reason why regenerative engineered vascular grafts are seen to be superior to engineered synthetic grafts ^10,11,19^. Two clinical trials have shown no infection in tissue-engineered vascular grafts^19^. However, little is known about how synthetic engineered or regenerative vascular grafts impact bacterial translocation. Although extensive work has been focused on the adhesion of bacteria to the surface of native^20^ and implanted materials^21^, none of this work has reduced infection to rates comparable or lower than blood vessels taken from the patient as autologous grafts^15,16,22,23^. Engineered vascular grafts made from expanded polyfluoroethylene (ePTFE) have been shown to prevent bacterial translocation in *in vitro* analysis^24^. However, Mufty et al., showed through meta-analysis that ePTFE vascular grafts and vascular grafts containing four antibiotics and silver as an antibacterial agent did not exhibit statistically significant levels of bacterial adhesion prevention or infection prevention either *in vitro* or in *in vivo* animal models ^16^.

Much of the focus in the literature on bacterial infection has been on adhesion to vascular graft materials specifically and medical implants materials in general^25-27^. Bacterial translocation describes the movement or passage of viable bacteria to normally sterile tissue or internal organs^28^. Surgical site infections, defined to be infection occurring within a 30-day window of surgery or within a year for implants, commonly result from bacterial translocation ^29^. While much focus has been given to sterile technique during surgery to reduce surgical site infections, microbiota have been found in almost every organ and tissue compartment in the human body^30-33^ potentially limiting the efficacy of better sterile technique. Bacterial translocation has been well studied in the gastrointestinal system and related surgeries ^34-36^. For example, *Escherichia coli* is the most frequently identified organism from gut translocation after laparotomy ^25,36,37^. Bacterial translocation can be rapid and has been shown to occur in the absence of systemic dissemination of bacteria in the blood. For example, *Staphylococci* translocate from the skin and from kidney infections within a time frame of 6 to 24 hours^38,39^. *Staphylococcus aureus* can cause deep tissue infections in the absence of an identifiable port of entry and can invade the bloodstream resulting in systemic bacterial infections^25,37^. While *E. coli* K1, *Streptococcus pneumonia*, and *Citrobacter spp*. do not lyse or otherwise damage microvascular endothelial cells when translocating from blood to the central nervous system across the blood-brain barrier ^40,41^. Although there is some correlation with bacteremia and translocation to various tissues, the concentration of blood-borne bacteria necessary to induce translocation in adult murine models is also enough to cause a variety of infections ^40^. Both the rapidity and immune evasion increase the challenges for diagnosis, treatment, and studies in whole-animal murine models ^42^.

Tissue engineered vascular grafts aim for the regeneration, repair, or buildup of functional vascular tissue which are functionally similar to the natural vessel ^43^. Tissue engineered vascular grafts utilizing host donor cells aim to reduce vascular graft fibrosis and rejection by the host immune system ^44-46^. It is hoped that this will lead to the usage of lowered levels of immunosuppressants, thereby improving the body’s overall ability to prevent bacterial infection ^17^. Hydrogels are frequently used for tissue engineering applications because they can cross-link and degrade *in situ*, eliminating the need for open surgery after the implanting process. Hydrogels like Gelatin Methacrylate (GelMA) and alginate are commonly used in tissue engineering due to their biocompatibility and the ease of tailoring their mechanical properties based on the application. 3D bio-printed hydrogel-based bio-ink provides a favorable environment for cell growth and tissue regeneration and has been effective in the regeneration of several types of tissue including heart, cartilage, muscle, kidney, and skin ^23,45,47-51^.

Decellularized tissue are similarly leveraged for tissue regeneration ^52^. Decellularized tissues utilizes organs or tissues from animals or lab-generated materials. Decellularization involves chemically or mechanically removing the living cells from a tissue, leaving behind a structured matrix material ^53-55^. Challenges of using decellularized tissues include promotion of host cell growth and differentiation, even though the latter is mediated by mechanical signaling similar to the tissue or graft desired^56,57^. Similar to engineered scaffolds populated with host cells, decellularized tissues are repopulated with host cells and are designed to minimize immune system rejection. However, bacterial infiltration into these matrices remains a concern ^52,58^. Hence, it is important to understand how bacteria translocate in decellularized tissue matrices and to highlight the potential challenges associated with infections in decellularized tissues.

This study aims to provide insight into the relationship between bacterial species and biomaterials by exploring the translocation of two common bacteria, *E. coli* and *S. aureus*, in alginate, GelMA, and decellularized porcine artery tissue. This study aims understand the viability of bacteria in these biomaterials over time and the bacteria’s potential for propagation or breakthrough out of the tissues, potentially elucidating some of the late stage (>30 days post operation) diagnosis of bacterial infection. Since *S. aureus* is known to cause deep tissue infection, we hypothesized that *S. aureus* would penetrate tissue better than *E. coli* while there would be no difference between *S. aureus* and *E. coli* penetration of engineered vascular graft materials GelMa and alginate.

## Experimental Methods

This study aims to provide insight into the relationship between bacterial species and biomaterials by exploring the translocation of two common bacteria, *E. coli* and *S. aureus*, in alginate, GelMA, and decellularized porcine artery tissue. The materials are prepared in three distinct methods. The constructs are then inoculated with the two different species at similar OD but differing concentration to optimize for visualization. The media from the midline of the construct is collected and plated to determine breakthrough. The infected constructs are then imaged using confocal microscopy. The process is outlined in Figure 1.

**Figure 1.**
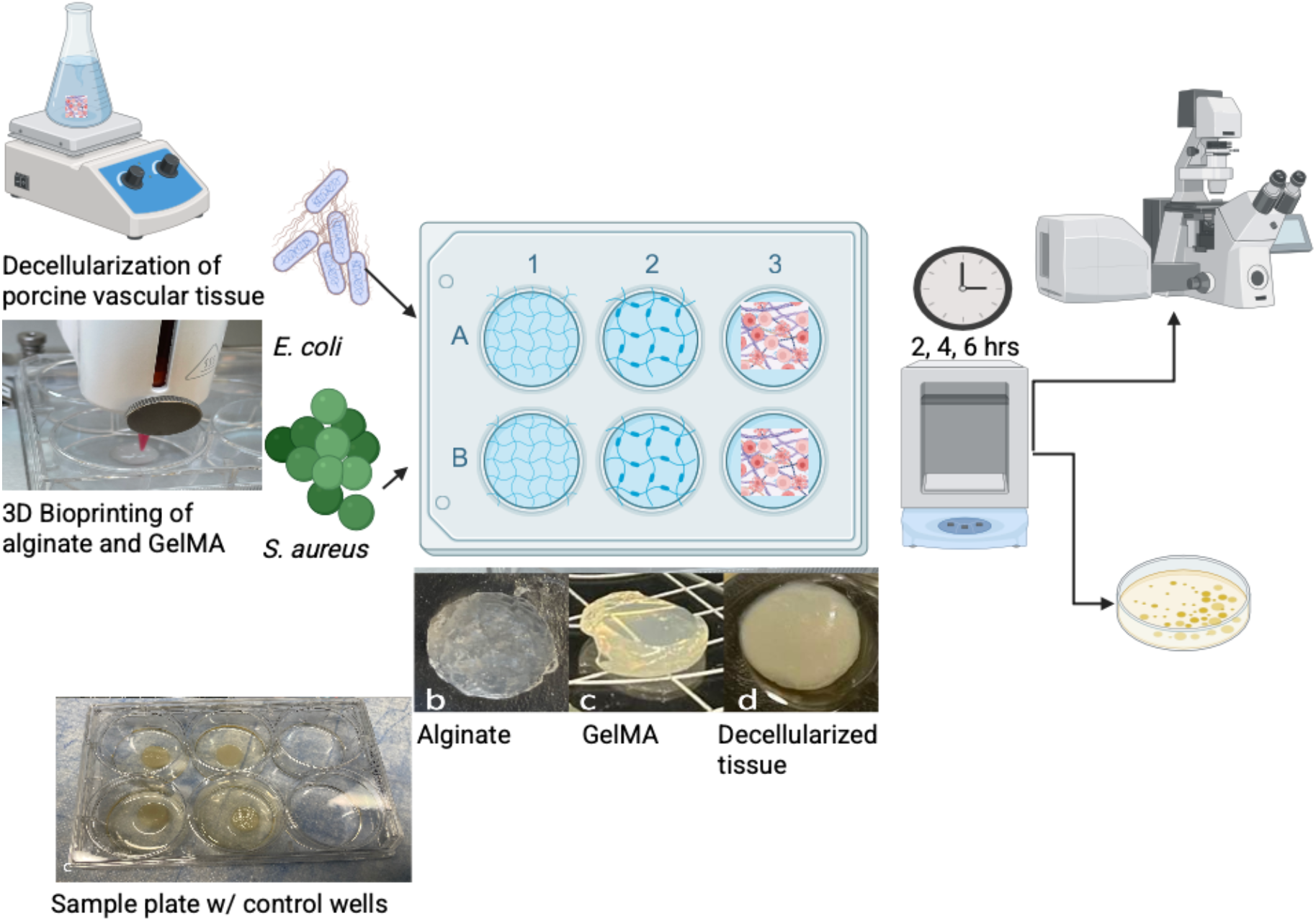
The process of decellularization of tissue and 3D bioprinting followed by inoculation of two bacterial species in three (b Alginate, c GelMA, d decellularized tissue) different tissue engineered constructs.

### Material printing, preparation, and sterilization

This study aims to understand the diffusion behavior of different types of bacterial species through different types of commonly used tissue engineering material (alginate and Gelatin Methacrylate hydrogels)^44,51,59^ relative to decellularized tissue (porcine blood vessels).

A 2% alginate construct was made by mixing 2% sodium alginate powder in distilled PBS and letting it sit in a dialysis tube inside calcium chloride for 24 hours. It was then cut into 3 mm OD by 3 mm thick samples using a die cutter for ease of handling. A Gelatin Methacrylate (GelMA) construct was created with the same dimensions (3 mm OD by 3 mm thick to be consistent) using the 3D extrusion printing (20 kPa, 2 mm/s speed and infill density at 100%, Cellink, Bico Group, GmBH, SWE) in a honeycomb pattern. The engineered materials were sterilized with UV light for two hours.

Three sets of porcine posterior vena cava were harvested from 10-12 days old Yorkshire pigs from North Carolina State University’s large animal laboratory within the post-mortem time of two hours (IACUC protocol number (21-205) from NCSU). While we cut the tissues to the same 3 mm diameter, we did not measure the thickness. Literature values indicate the vena cava of young pigs is 0.6 mm ^60^. The extracted blood vessels were immediately washed with phosphate-buffered saline (PBS) solution to remove excess blood and debris. The cleaning process was followed by a decellularization process by: treating the blood vessels with sodium dodecyl sulfate (SDS) solution and penicillin-streptomycin for 48 hours and washing with 1x PBS for 24 hours in an orbital shaker ^61^. The decellularized tissue was sterilized using 2% peracetic acid with 100% ethanol and distilled water in the ratio of 2:1:1 and rinsed three times alternating between ethanol and sterile saline solution for 15 minutes. Penicillin oxidizes in peracetic acid ^62^ while streptomycin is likely degraded by peracetic acid ^63^. Decellularization was verified using a DNeasy (QIAGEN®, Germany) kit and nanodrop UV5 Nano (Mettler Toledo). The nanodrop quantification was done for *n* = 3 animals for both decellularized and native tissues.

Alginate samples were made by casting, GelMA samples were made by 3D bioprinting, and tissue samples were made by decellularizing. After preparation, the samples were cut into small pieces. The frozen samples were put into a freeze dryer (lyophilizer) without letting them thaw. The freeze dryer was set to a vacuum pressure of 10-20 millitorr and a temperature below -40°C. The samples were left to dry completely over 24 hours. Once freeze-dried, the samples were carefully handled to avoid damage. The dried samples were mounted onto SEM stubs using carbon tape. The samples were coated with a thin layer of gold using a sputter coater. The mounted samples were placed in SEM for imaging.

### Bacteria Inoculation and Seeding

The guidelines for testing vascular graft tissue engineered medical products does not include recommendations for infection testing ^64^. Previously, guidance on hierarchical integration did not include recommendations for timing or inoculation ^65^. Studies on *in vivo* infections have ranges from 10 – 10^5^ CFU/mL ^58,66^

*Escherichia coli* (ATCC 12435) and *Staphylococcus aureus* (ATCC 25923) bacteria were inoculated in BD Difco™ Ampicillin (AMP) and Tryptic Soy Broth (TSB), respectively. *E. coli* (ATCC 12435) exhibits low levels of streptomycin and ampicillin resistance while *S. aureus* (ATCC 25923) is a susceptible reference strain. *E. coli* underwent an overnight incubation in AMP media for eight hours, while *S. aureus* was cultured in TSB. Following the eight-hour incubation period, both bacterial cultures were standardized to an optical density of 0.2 at 600 nm, which yielded bacterial concentrations of 9.95×10^6^ CFU/mL for *E. coli* and 2.15×10^6^ CFU/mL for *S. aureus*. This concentration is likely higher than what is introduced by non-sterile techniques, it is the lowest concentration we found that would survive the experiment and be visualizable in the construct of the selected sizes.

Both *E. coli* and *S. aureus* bacteria were incubated within two sets of polymeric materials (GelMA 3D constructs and alginate hydrogels) and one set of biological materials (decellularized porcine vessel). The translocation of each type of bacteria was monitored at precise time intervals of two, four, and six hours of post-inoculation in each of the materials. We selected six hour maximum time to minimize bacteria growth in the construct while further acknowledging that the after six hours innate immune response would affect the *in vivo* results. We further selected these time points to minimize tissue utilization while maximizing the temporal resolution. In the case of GelMA, decellularized tissue, and alginate material, 10 µL of each bacterial suspension was introduced onto the top of respective substrates. Media was placed up to the midline of the construct, simulating the more nutrient rich blood stream relative to the tissue environment, in this case a material-air interface. After each time point, the media was removed surrounding the constructs, and CFU/mL counts were generated for all replicates. One well without constructs but with media was inoculated with bacteria as a positive growth control. The materials were then subjected to a thorough sterile saline wash to eliminate any unattached bacteria and imaged with confocal microscopy.

### Image Analysis and cell count

After each time point (2, 4, 6 hrs), bacteria-seeded constructs, i.e. GelMA, alginate, and decellularized porcine tissue, were incubated in staining media supplemented with 4 mmol/L calcein AM and 2 mmol/L ethidium-homodimer for 20 min at 37°C. Each of the constructs were then washed with PBS solution and covered with glass coverslips for conducting confocal microscopic analysis. Confocal fluorescence microscopy (Zeiss LSM 900, Zeiss GmBH) was used for taking high-resolution 3D images and the bacterial count was analyzed at three time points (*n*=3 per construct, per biological replicate, per time point) using ImageJ software (National Institutes of Health).

### Mid-point Analysis and Statistical Analysis

The ability of both *E. coli* and *S. aureus* bacteria to penetrate all three constructs was quantified using midpoint analysis. Bacterial cell counts were measured via z-stacks at volumes within the construct for each time. The cell counts were per slice. The midpoint and weighted centroid of the cell construct were computed. Trials were coincident, so we computed penetration via midpoint analysis for speed.

All experimental results were expressed in the form of mean ± standard deviation. The Shapiro-wilk test was conducted to examine the normality test. A multiway ANOVA followed by a posthoc test for multiple pairwise comparisons was performed to evaluate statistical differences between the GelMA, alginate, and decellularized porcine vessel, at three time points as described in Table 1 and Table 2. A value of p< 0.05 is assumed to be statistically significant in this study.

**Table 1.** Breakthrough aggregated number of samples.

| Hour | Aggregated number of samples |
| --- | --- |
| 2 | 24 |
| 4 | 24 |
| 6 | 24 |
| Material | n |
| Alginate | 18 |
| Gelma | 18 |
| PC | 18 |
| Tissue | 18 |
| Species | n |
| *E. coli* | 36 |
| *S. aureus* | 36 |

**Table 2.** ANOVA table for breakthrough comparing Species (*E. coli* and *S. aureus*), Material (GelMA, alginate, decellularized tissue, positive control), and Hour (2 hours, 4 hours and 6 hours) Df: Degrees of freedom; Sum Sq: Sum of Squares; Mean Sq: Mean Square Error; F value; P-value.

|  | Df | Sum Sq | Mean Sq | F value | P-value |
| --- | --- | --- | --- | --- | --- |
| Species | 1 | 0.54 | 0.544 | 0.347 | 0.558813 |
| Material | 3 | 91.37 | 30.458 | 19.42 | 2.20E-8 |
| Hour | 2 | 8.69 | 4.346 | 2.771 | 0.072649 |
| Species : Material | 3 | 39.27 | 13.09 | 8.347 | 0.000144 |
| Species : Hour | 2 | 3.01 | 1.504 | 0.959 | 0.390441 |
| Material : Hour | 6 | 17.03 | 2.838 | 1.81 | 0.117138 |
| Species : Material: Hour | 6 | 10.93 | 1.821 | 1.161 | 0.342543 |

## Results

The goal of this study was to determine how bacteria penetrate and breakthrough tissue engineered materials used in vascular grafts relative to decellularized vascular tissue. This was studied using SEM, tensile testing, confocal microscopy, and CFU counts of two engineered materials, GelMA and alginate, and decellularized porcine vascular tissue. Materials were characterized using SEM and tensile testing. Breakthrough of bacteria introduced on top of the material was analyzed using confocal microscopy and colony forming units.

Increased porosity of vascular graft material should increase bacterial translocation if the pore sizes are greater than the diameter of the bacterial cells. SEM of the surface of alginate, 3D printed GelMA, and decellularized vascular tissue showed no discernable differences, Figure 2. We did not take cross sections of the material, however, cracks in the alginate showed potential pore structures < 5 µm in diameter.

**Figure 2.**
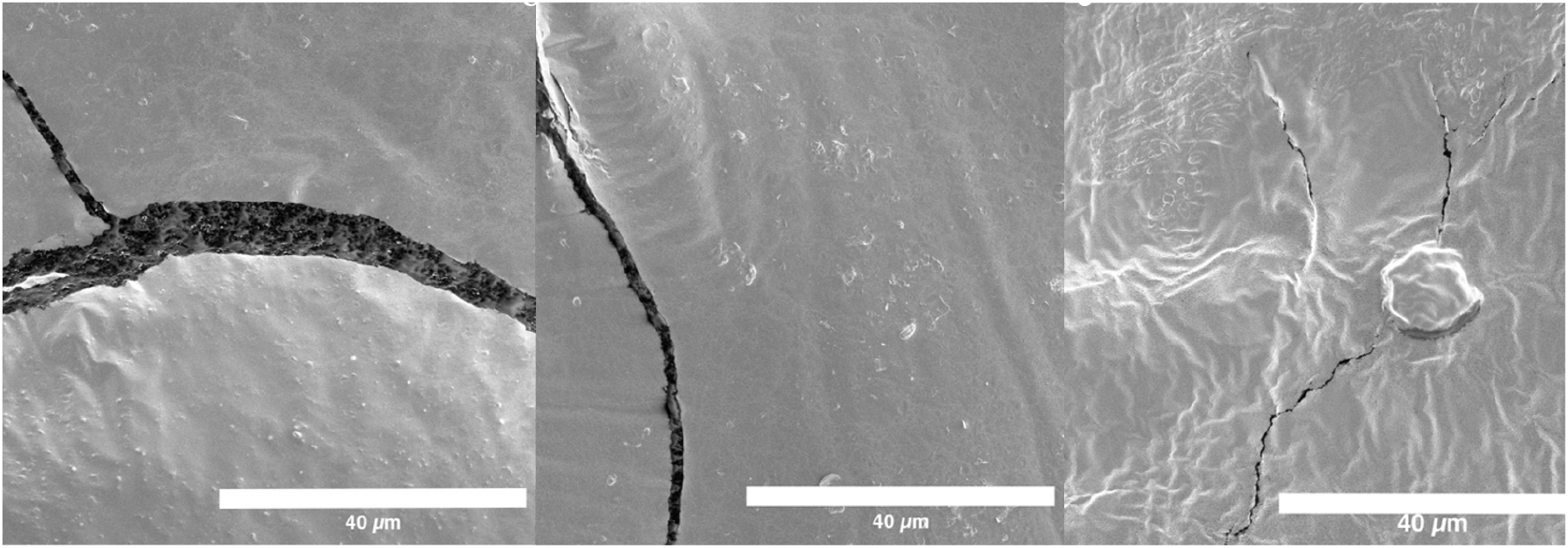
SEM Images of (a) alginate, (b) GelMA and (c) decellularized vascular tissue GelMA

To measure bacterial translocation, confocal microscopy was used at 2, 4, and 6 hrs after seeding. Figure 3 and Figure 4, show representative 3D views of GelMA, alginate and decellularized tissue constructs after 2,4 and 6 hrs of seeding with *S. aureus* and *E. coli* bacteria respectively (all images available in the Duke Research Data Repository^67^). Live-dead stains were used to indicate viability of the bacterial cells inside the construct. While some of the chemicals used in decellularization may lead to cell death, there was no indication this was occurring, Figure 3 and Figure 4. *S. aureus* ATCC 25923 as a reference strain is susceptible to both antibiotics used in the decellularization process ^68^, while there is limited information on *E. coli*. We did not observe any bacteria in the negative control. These images were then used to measure the extent of bacterial cell penetration.

**Figure 3.**
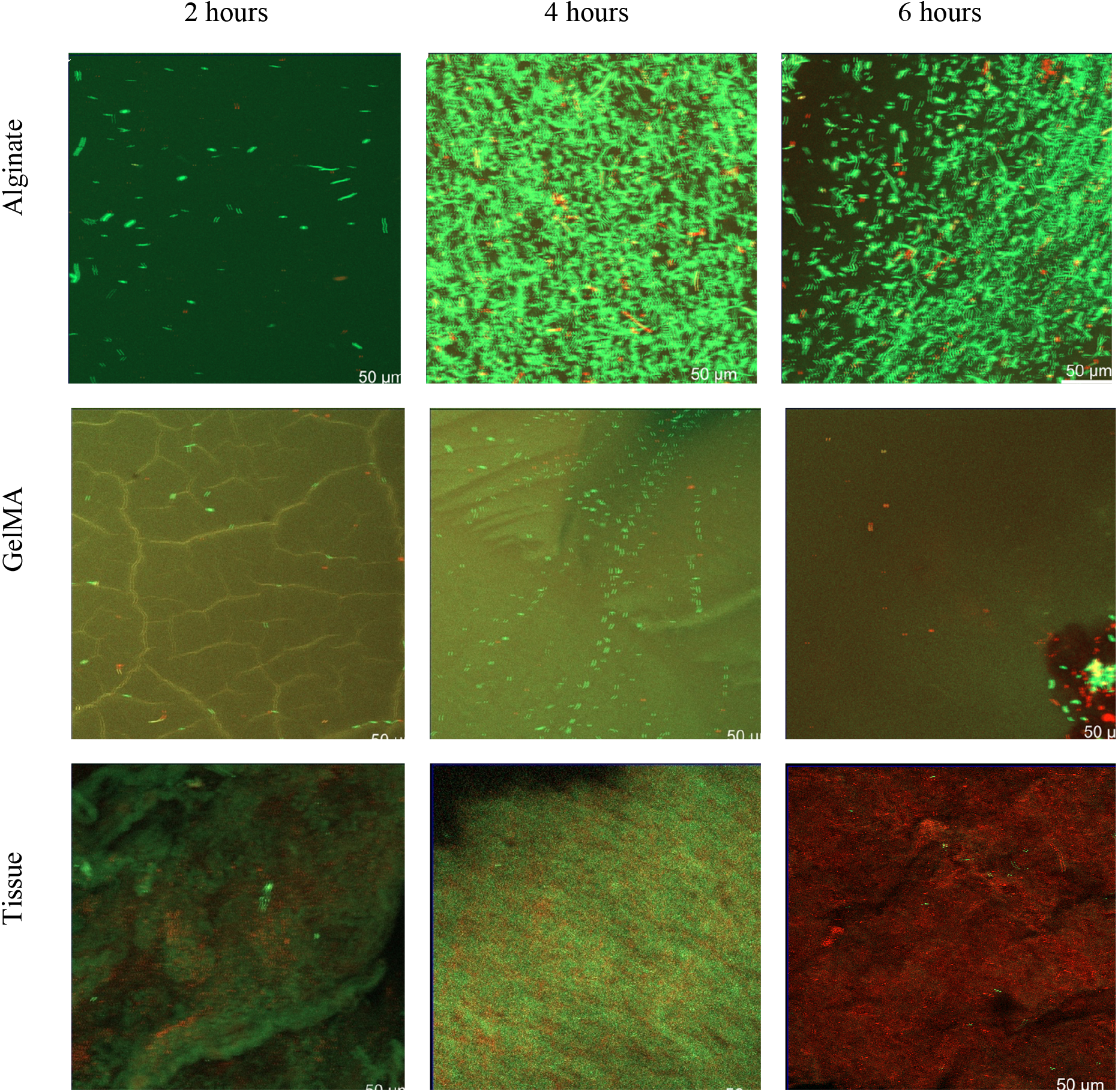
Representative confocal microscopy images of *E. coi* in alginate (a-c) at 2, 4, 6 hours respectively. Representative confocal microscopy images of *E. coli* in GelMA (d-f) at 2, 4, 6 hours respectively. Representative confocal microscopy images of *E. coli* in tissue samples at 2, 4, 6 hours respectively. These images are the Z-plane of the maximum intensity projections of 3D stacks of the samples onto a 2D plane, red are dead cells, green are live cells.

**Figure 4.**
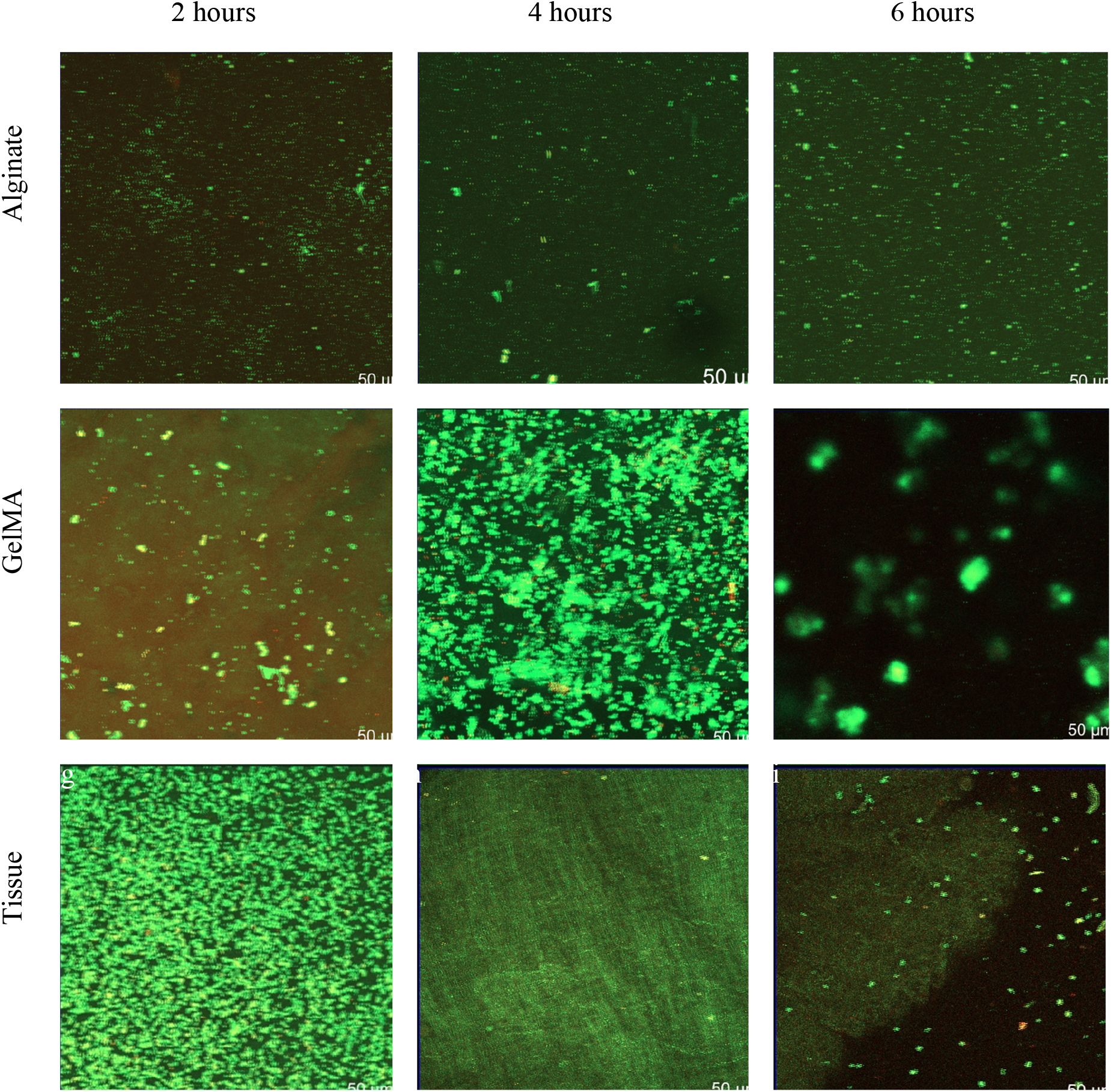
Representative confocal microscopy images of *S. aureus* in alginate (a-c) at 2, 4, 6 hours respectively. Representative confocal microscopy images of *S. aureus* in GelMA (d-f) at 2, 4, 6 hours respectively. Representative confocal microscopy images of *S. aureus* in tissue samples at 2, 4, 6 hours respectively. These images are the Z-plane of the maximum intensity projections of 3D stacks of the samples onto a 2D plane, red are dead cells, green are live cells.

There was consistent stage jitter artifacts in the images, notably for *E. coli* in GelMA as seen in Figure 3e. Since penetration is calculated from the centroid of the mass, these artifacts cancel out during computation of penetration depth. Also of note is the GelMA porous structure can be seen and appears to constrain the *E. coli* at the 2 hr mark, Figure 3d. While this constraint was not seen in subsequent time points (4 and 6 hrs), this expected constraint on bacterial cell migration needs to be further studied to understand its potential impact on infection. While it may be reasonable to conclude from Figure 3f-g that bacteria cells localized to tissue, we did not stain tissue specifically. Future work optically profiling tissues is needed to understand if this is localization.

As expected, there was more clustering among *S. aureus* relative to *E. coli* as seen in Figure 4d-f. GelMA structures were not visible in the images unlike in the *E. coli* images. Tissue structures were similarly not as visible in the images as in *E. coli*.

Bacteria species appears to have a statistically significant effect on the penetration depth into the construct with *E. coli* reaching *∼*1.5 mm on average while *S. aureus* reached ∼2 mm in depth, Figure 5 (data in SI Table 1). The material of the construct did not appear to have a statistically significant effect on penetration depth. Similarly, time did not appear to have a statistically significant effect on penetration depth.

**Figure 5.**
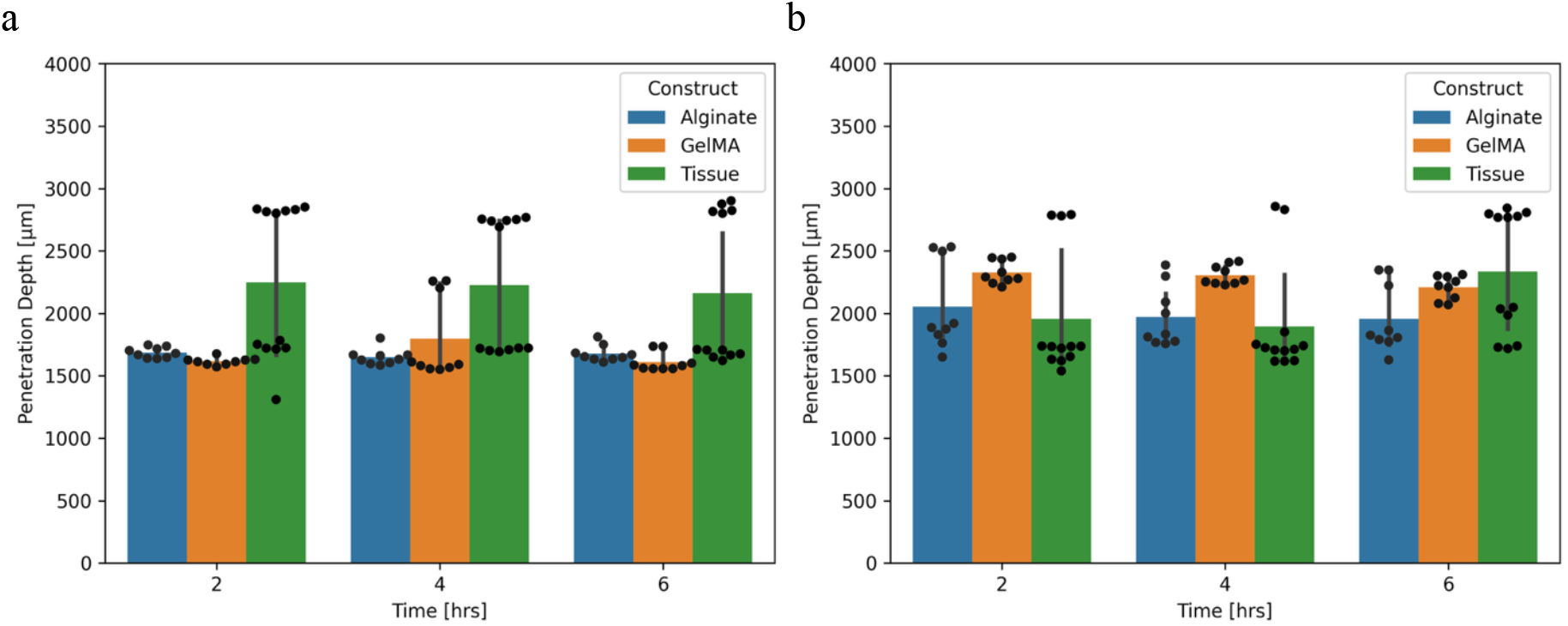
a. Average *E. coli* penetration into alginate, GelMA, and porcine tissue. b. Average *S. aureus* penetration depth into alginate, GelMA, and porcine tissue. Data points shown includes 3 technical replicates on 3 biological replicates. Mean and 95% confidence interval statistics shown for averages of technical replicates with bootstrapping.

As shown in Figure 6, *E. coli* and *S. aureus* breakthrough about the same for tissue, alginate, GelMA and the positive control initially (data in SI Table 2). After 4 hours, there is a statistically significant difference (p<0.05) for *E. coli*, showing decreased proliferation in tissue relative alginate, GelMA and the positive control after 6 hours. After 4 hours there is a statistically significant difference (p<0.05) for *S. aureus* showing increased breakthrough in GelMA relative to the positive control. After 6 hours, there is a statistically significant difference (p<0.05) increase in breakthrough of *S. aureus* relative to decellularized tissue. As described in Table 2, multiway ANOVA however, showed temporal differences but differences between materials and differences between species-material parings.

**Figure 6.**
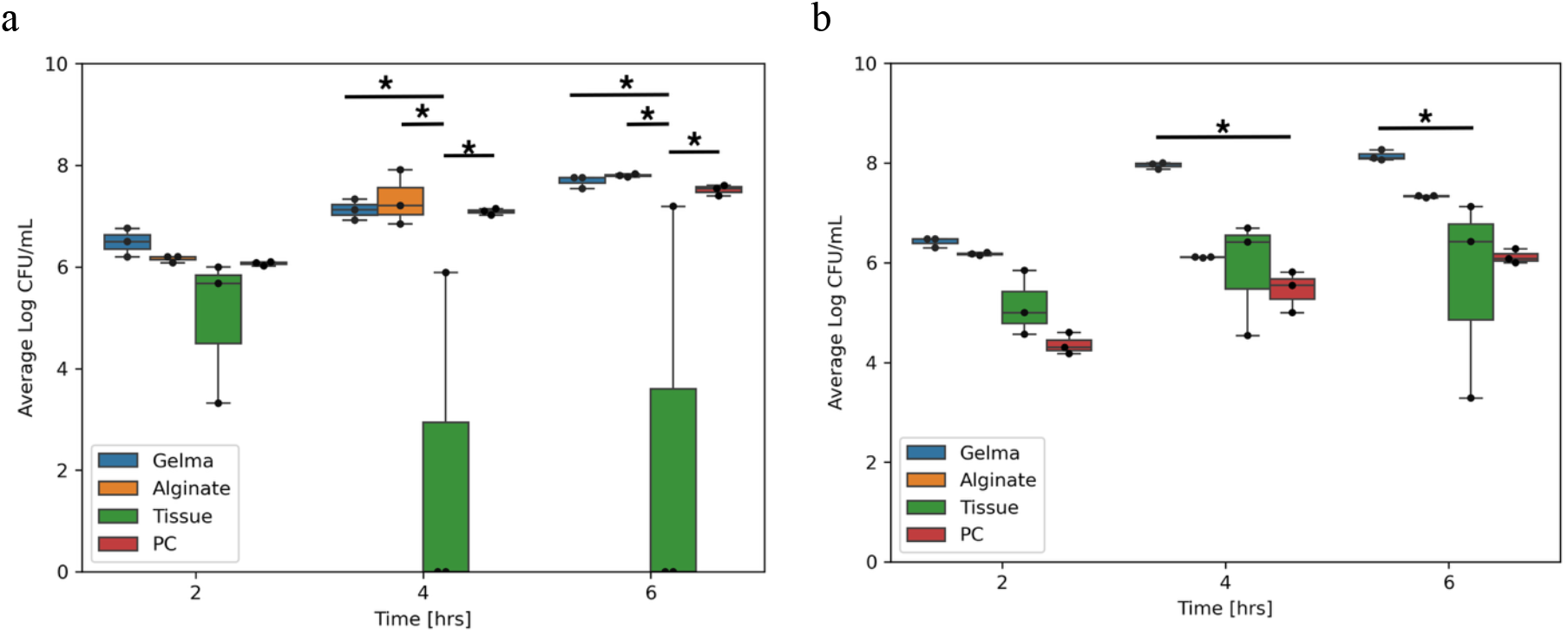
a. Breakthrough rates represented by averages of three technical replicates for three biological replicates in Log CFU/mL of *E. coli* and b. *S. aureus*. One-way ANOVA Tukey multiple comparisons of means 95% family-wise confidence level. (* p<0.05, ** p<0.01, *** p<0.001, **** p<0.0001)

## Discussion

The goal of this study was to quantify bacterial penetration and breakthrough differences between tissue engineered materials, GelMA, alginate, and decellularized porcine vascular tissue. While previous studies had shown that there was no penetration or breakthrough in synthetic vascular grafts made from expanded polyfluoroethylene (ePTFE) and polyurethane (PU) there has not been a study on penetration rates and proliferation in tissue engineered materials and decellularized tissue. We observed no differences in bacteria penetration depth between species or between constructs. We observed reduction in breakthrough in decellularized tissue relative to all constructs for *E. coli* while *S. aureus* had increased levels of breakthrough in GelMA relative to the positive control and tissue at 4 and 6 hours respectively.

This study focused on penetration into a lumen-like space, a reservoir of media at the midline of the construct. The midpoint of both species penetration reached the midline, ∼1.5 mm, with *S. aureus* reaching deeper than *E. coli*. Species dependent penetration parameters include species size, motility, anchoring strategies, chemotactic receptors, and quorum sensing receptors. *S. aureus* and *E. coli* have similar minor diameters, ∼ 1 µm. *S. aureus* is approximately spheroidal and *E. coli* is rod-shaped ranging from 2 – 8 µm in length depending on the stress conditions ^69^. *S. aureus* and *E. coli* have distinct modes of motility ^70^. *S. aureus* moves primarily through sliding a diffusion-like process, sometimes a ‘comet’ gliding like motility ^70,71^ while *E. coli* has swimming motility via run and tumble ^72^. Sliding motility is strictly slower than swimming. The observed pore size, < 5 µm, may have reduced the speed of swimming motility of *E. coli* since run and tumble exposes the major axis ^72^.

The absence of measured difference in penetration speed, that is depth over time, may be attributable to the sampling frequency or limited duration of our experiments. For example, traveling bands of *E. coli* ^73,74^ have been observed at 6 hours, the maximum duration of the experiment in this study. The limited duration was intentional in this study design to not overlap with *in vivo* cellularization of tissue engineered scaffolds. Cellularization occurs after 24 hours and confers immunity dependent on the engineered tissue design and host factors beyond fundamental interactions of material and bacteria ^26,65^. While a limitation, similar penetration between species is consistent with the non-specificity of bacterial species infections in literature

^18^. Future work increasing sampling frequency may show more differences in species penetration. The mechanism of penetration depth differences may have been further elucidated with staining and imaging of the material constructs as there was visible constriction in Figure 3d that was not able to be visualized in any other figure. *S. aureus* anchors to the von Willebrand Factor using *Staphylococcus* protein A in native vasculature which may pull it through pores ^75^. This factor is not present in engineered or decellularized tissue. Similarly, flagella mediated wetting, where rotating flagella of *E. coli* push water into pores, may have offset the size effect. Similarly, the production of surfactants may have increased the speed of *S. aureus* ^76^. *E. coli* leverages both chemotaxis and quorum sensing strategies ^77^, while *S. aureus* only leverages quorum sensing ^70^. Future work could address these factors using mutant strains that lack flagella, chemotactic pathways, and quorum sensing pathways. Furthermore, quorum sensing may have skewed the results in favor of this midline position, as those chemicals could freely diffuse once the midpoint was reach.

We showed decreased breakthrough in decellularized tissue relative to engineered materials for both species. Multiway ANOVA, grouping by material including all time points and all species, there are 18 samples per material leading to high statistical power in post-hoc analysis. This difference was more pronounced for *E. coli* than for *S. aureus*, however with less confidence. Furthermore, two out of three biological replicates of *E. coli* in decellularized tissue showed no measurable breakthrough. *E. coli* uses both chemotaxis and quorum sensing while *S. aureus* only uses quorum sensing. The differences in breakthrough may be attributable to concentration gradients in the engineered material being larger than the decellularized material. Another potential cause of differences breakthrough may be connected to heterogeneity. A common hypothesis is spatial heterogeneity of chemical and structural components is important engineered tissue regeneration. While it is well known that hydrogels are homogeneous. Our findings suggest that heterogeneity may also be important for infection prevention.

The absence of intraspecies effects, one-way ANOVA on material for fixed time points either *E. coli* or *S. aureus*, is likely a function of sample size limiting statistical power. However, the rare breakthrough of *E. coli* is supported the rare reports of vascular graft infection ^18,19^. Further work should be done over longer times, more and thicker decellularized vascular tissue types.

Our findings suggest that heterogeneous materials may have antibacterial impacts. Heterogeneous materials have been suggested to improve function and cellularization ^23,44,46,49^. This heterogeneity was marginally visible between the outer surface and cracks in the SEM images, Figure 2. It was more visible in confocal images, noting the well mixed appearance of bacteria in hydrogels relative to structure localized bacteria in decellularized vasculature, Figure 3 and Figure 4. The delayed infection rates of vascular grafts may be due to penetration of bacteria being independent of species and material while the breakthrough rates were dependent. This would leave a source of bacteria for infection after antibiotics have been stopped post-surgery. Since most vascular grafts are not made from tissue engineered materials, this homogeneity would necessarily come from sutures or from host coating the vascular graft. Sutures have been shown to produce acellular regions that may be more homogeneous ^78^. Future work should be performed on decellularized *ex vivo* tissues post-surgery to determine how these structures may enable bacteria proliferation.

## Conclusions

Vascular grafting is a key component of medical care that millions use annually. While autologous vascular grafts are superior, there are often not enough or not the right size. Instead, engineered vascular grafts must be used. In rare, but often fatal, cases, engineered vascular grafts get infected. While much of the work on vascular graft infection has focused on adhesion, there has been little study of bacterial penetration and breakthrough. Here we examined penetration of *E. coli* and *S. aureus* into GelMa and alginate and decellularized porcine vascular tissue. We showed similar penetration rates and penetration depths, differing by material but not species. We found species dependent differences in breakthrough. Specifically, we found decreased breakthrough of *E. coli* from decellularized tissue relative to engineered materials. As the penetration and penetration rates were similar, the lack of breakthrough may contribute to the delayed onset of infection seen in the literature. The difference between hydrogels and decellularized tissue is homogeneous structure instead of heterogeneous structure. We propose that heterogeneous structures may reduce infection of tissue engineered vascular grafts.

## Supporting information

Supplemental Experimental Figures and Data tables

## Acknowledgments

Research reported in this publication was supported by the National Institute of General Medical Sciences of the National Institutes of Health under Award Number NIH R35 GM142898. The content is solely the responsibility of the authors and does not necessarily represent the official views of the National Institutes of Health. The authors thank Tim Boston at North Carolina State University for porcine vascular samples. The authors would also like to thank Fredrick Ausubel at Harvard University for his feedback and discussions on early drafts of the manuscript.

## Author contributions

A.S. and A.J. designed the experiment. A.S. performed the experiment. J.P. performed data analysis. A.S. wrote the original draft. A.S., J.P., and A.J. reviewed and edited the draft. A.J. conceived, administered and supervised the project.

## Data Availability

The data that support the findings of this study are openly available in Duke Research Data Repository at https://doi.org/10.7924/r4v98dn1x, reference number 67.

