## Supplemental Experimental Figures and Data tables for "*In vitro* evaluation of *Escherichia coli* and *Staphylococcus aureus* translocation in 3D printed material"

Authors: Ashma Sharma[1a], Joshua Prince[1b], A-Andrew D Jones III[1234c*]

Affiliations:

1. Department of Civil & Environmental Engineering, Pratt School of Engineering, Duke University, Durham, NC 27708
2. Thomas Lord Department of Mechanical Engineering & Materials Science, Pratt School of Engineering, Duke University, Durham, NC 27708
3. Duke Microbiome Initiative, Duke University, Durham, NC 27708
4. Duke Materials Initiative, Duke University, Durham, NC 27708
    b. Present address: ICF International ORCID: 0000-0001-6050-9544

*Corresponding Author

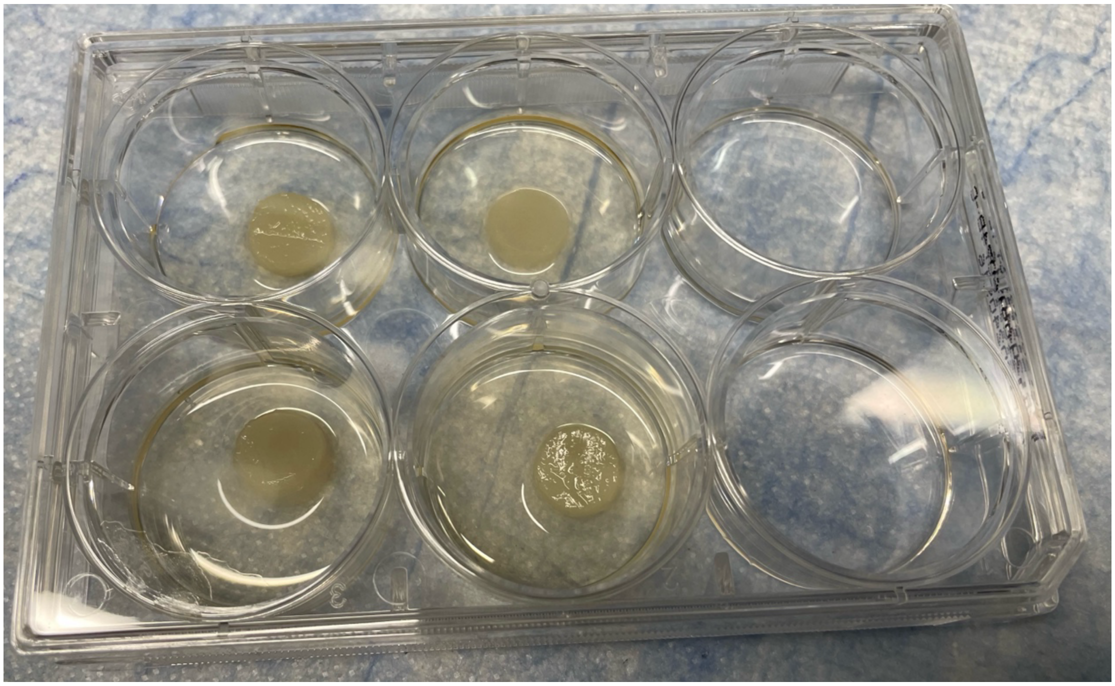

SI Figure 1: Culturing of bacteria against construct in 6-well plates. The media is filled to the midline of the biomaterial construct.

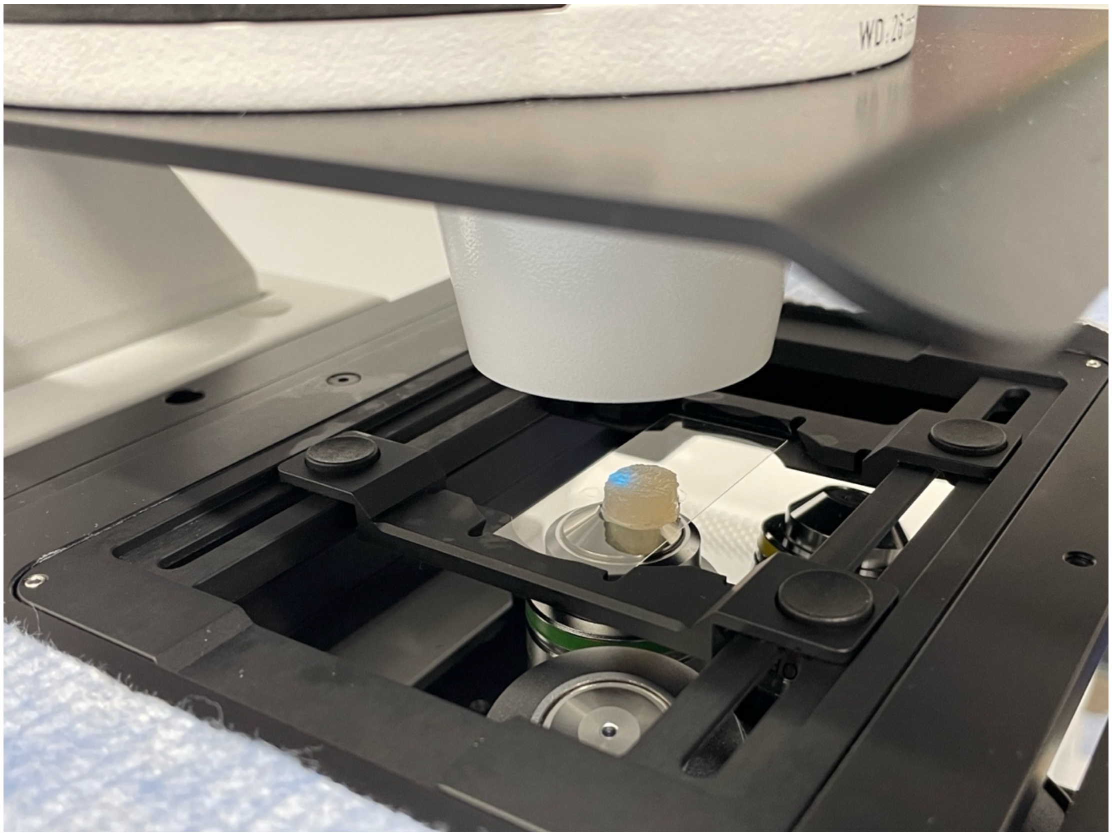

SI Figure 2. Set up of the confocal microscopy of each construct. Glass slides were used to balance the construct and images were taken using an inverted microscope.

**SI Table S1.** ***E. coli*** **and** ***S. aureus*** **penetration into alginate, GelMA, and porcine tissue mean and 95% confidence interval statistics shown for averages of technical replicates with bootstrapping from Figure 5.**

| Material | Species | Biological Replicate Average | Biological Replicate SD | Biological Replicate 95%CI Magnitude | Time [hrs] |
|---|---|---|---|---|---|
| Tissue | *E. coli* | 2246.375 | 368.910 | 361.526 | 2 |
| Tissue | *E. coli* | 2226.417 | 37.659 | 36.905 | 4 |
| Tissue | *E. coli* | 2162.875 | 641.807 | 628.959 | 6 |
| GelMA | *E. coli* | 1615.667 | 54.212 | 61.346 | 2 |
| GelMA | *E. coli* | 1797.667 | 32.194 | 36.431 | 4 |
| GelMA | *E. coli* | 1607.778 | 84.555 | 95.681 | 6 |
| Alginate | *E. coli* | 1685.778 | 43.900 | 49.677 | 2 |
| Alginate | *E. coli* | 1648.889 | 118.140 | 133.686 | 4 |
| Alginate | *E. coli* | 1676.556 | 67.841 | 76.768 | 6 |
| Tissue | *S. aureus* | 1955.667 | 135.179 | 132.473 | 2 |

| | | | | | |
|---|---|---|---|---|---|
| Tissue | *S. aureus* | 1894.083 | 432.366 | 423.711 | 4 |
| Tissue | *S. aureus* | 2334.667 | 45.352 | 44.444 | 6 |
| GelMA | *S. aureus* | 2327.744 | 53.558 | 60.605 | 2 |
| GelMA | *S. aureus* | 2306.778 | 105.998 | 119.946 | 4 |
| GelMA | *S. aureus* | 2207.111 | 61.800 | 69.932 | 6 |
| Alginate | *S. aureus* | 2053.011 | 110.957 | 125.558 | 2 |
| Alginate | *S. aureus* | 1969.000 | 363.284 | 411.086 | 4 |
| Alginate | *S. aureus* | 1955.667 | 141.720 | 160.368 | 6 |

**SI Table 2. Breakthrough rates represented by averages of three technical replicates for three biological replicates in Log CFU/mL of *E. coli* and *S. aureus* from Figure 6. One-way ANOVA Tukey multiple comparisons of means 95% family-wise confidence level.**

| Material | Species | Biological Replicate Average | Biological Replicate SD | Biological Replicate 95%CI Magnitude | Time [hrs] |
|---|---|---|---|---|---|
| Tissue | *E. coli* | 4.996 | 1.462 | 3.632 | 2 |
| Tissue | *E. coli* | 1.963 | 3.400 | 8.446 | 4 |
| Tissue | *E. coli* | 2.398 | 4.153 | 10.316 | 6 |
| PC | *E. coli* | 6.066 | 0.040 | 0.099 | 2 |
| PC | *E. coli* | 7.088 | 0.063 | 0.156 | 4 |
| PC | *E. coli* | 7.515 | 0.105 | 0.261 | 6 |
| GelMA | *E. coli* | 6.486 | 0.282 | 0.701 | 2 |
| GelMA | *E. coli* | 7.125 | 0.207 | 0.513 | 4 |
| GelMA | *E. coli* | 7.685 | 0.124 | 0.309 | 6 |
| Alginate | *E. coli* | 6.160 | 0.055 | 0.136 | 2 |
| Alginate | *E. coli* | 7.321 | 0.543 | 1.348 | 4 |
| Alginate | *E. coli* | 7.799 | 0.030 | 0.075 | 6 |
| Tissue | *S. aureus* | 5.138 | 0.652 | 1.618 | 2 |
| Tissue | *S. aureus* | 5.882 | 1.171 | 2.909 | 4 |
| Tissue | *S. aureus* | 5.612 | 2.044 | 5.078 | 6 |
| PC | *S. aureus* | 4.360 | 0.219 | 0.544 | 2 |
| PC | *S. aureus* | 5.452 | 0.414 | 1.029 | 4 |
| PC | *S. aureus* | 6.119 | 0.144 | 0.357 | 6 |
| GelMA | *S. aureus* | 6.418 | 0.102 | 0.252 | 2 |
| GelMA | *S. aureus* | 7.951 | 0.067 | 0.166 | 4 |
| GelMA | *S. aureus* | 8.142 | 0.110 | 0.273 | 6 |
| Alginate | *S. aureus* | 6.175 | 0.029 | 0.072 | 2 |
| Alginate | *S. aureus* | 6.108 | 0.010 | 0.024 | 4 |

| | | | | | |
|---|---|---|---|---|---|
| Alginate | *S. aureus* | 7.328 | 0.024 | 0.059 | 6 |
